# Reversion to sensitivity explains limited transmission of resistance in a hospital pathogen

**DOI:** 10.1101/2024.06.03.597162

**Authors:** Kevin C. Tracy, Jordan McKaig, Clare Kinnear, Jess Millar, Aaron A. King, Andrew F. Read, Robert J. Woods

## Abstract

Bacterial pathogens that are successful in hospital environments must survive times of intense antibiotic exposure and times of no antibiotic exposure. When these organisms are closely associated with human hosts, they must also transmit from one patient to another for the resistance to spread. The resulting evolutionary dynamics have, in some settings, led to rising levels of resistance in hospitals. Here, we focus on an important but understudied aspect of this dynamic: the loss of resistance when the resistant organisms evolve in environments where the antibiotic pressure is removed. Based on prior data, we hypothesize that resistance arising in the context of strong selection may carry a high cost and revert to sensitivity quickly once the selective pressure is removed. Conversely, resistant isolates that persist through times of no antibiotic pressure should carry a lower cost and revert less quickly. To test this hypothesis, we utilize a genetically diverse set of patient-derived, daptomycin-resistant *Enterococcus faecium* isolates that include cases of both *de novo* emergence of resistance within patients and putatively transmitted resistance. Both of these sets of strains have survived periods of antibiotic exposure, but only putatively transmitted resistant strains have survived extended periods without antibiotic exposure. These strains were then allowed to evolve in antibiotic free laboratory conditions. We find that putatively transmitted resistant strains tended to have lower level resistance but that evolution in antibiotic-free conditions resulted in minimal loss of resistance. In contrast, resistance that arose *de novo* within patients was higher level but exhibited greater declines in resistance *in vitro*. Sequencing of the experimentally evolved isolates revealed that reversal of high level resistance resulted from evolutionary pathways that were frequently genetically associated with the unique resistance mutations of that strain. Thus, the rapid reversal of high-level resistance was associated with accessible evolutionary pathways where an increase in fitness is associated with decreased resistance. We describe how this rapid loss of resistance may limit the spread of resistance within the hospital and shape the diversity of resistance phenotypes across patients.

## 2 Introduction

Antibiotic resistance is a significant public health concern, particularly in hospitals, where increasing levels of resistance are facilitated by two processes that occur at elevated rates. First, strong selective pressure from therapeutic usage of antibiotics acts to promote the *de novo* evolution of resistance within hosts (*1, 2*). Second, hospitals function as hot spots for transmission, where large numbers of contacts between patients increase the spread of isolates (*3*). Despite the selection and transmission of resistant isolates, antibiotic resistance often does not reach universally high levels (*4*). This constraint on the evolution of resistance is rarely due to the inability of the organisms to evolve resistance: It is frequently possible to evolve high-level resistance in the laboratory (*5, 6*). Rather, resistance evolution appears constrained by pleiotropic effects on the ability to survive within the patient and transmit to other patients in environments without antibiotics (*7*).

In this work, we seek to understand these constraints on resistance evolution by investigating the evolutionary steps that lead to the successful transmission of resistant mutants. We study the nosocomial pathogen *Enterococcus faecium* and the evolution of resistance to the antibiotic daptomycin, which remains a preferred treatment for these infections (*8*). Daptomycin resistance was rare in *E. faecium* before the widespread use of daptomycin, but has increased concurrently with daptomycin usage (*4, 9, 10*). Resistance evolution within patients (*11, 12*) and transmission between patients have been well documented (*13, 14*). Thus, daptomycin resistance in *E. faecium* represents a clinical setting where the early steps toward the evolution of transmissible resistance can be studied. Specifically, we hypothesize that robustness to phenotypic reversions in antibiotic-free conditions plays a significant role in the spread of nosocomial resistant pathogens.

The conceptual basis of our hypothesis is as follows. Resistance emerges when an antibiotic-sensitive bacterial population is exposed to an antibiotic. This newly resistant population must continue to transmit between hosts in antibiotic free conditions to ensure its survival. Multiple evolutionary processes may occur during the transmission process and with the removal of antibiotic selective pressure (*15*). First, evolution may lead to the loss of resistance, either through precise molecular reversions or secondary mutations that negate the impact of the initial resistance mutation. In this case, we would be unlikely to observe reversion and it would not contribute to the problem of transmitted resistance. Second, compensatory mutations may arise which maintain the resistance phenotype while alleviating the costs of resistance (*5, 16–18*).

Third, a subset of resistance mutations may have low or no fitness cost in antibiotic free conditions. In this case, repeated sampling of the mutational landscape may allow some populations to find these lower-cost resistance phenotypes. The latter mechanism would imply that *de novo* evolution produces a range of resistance mechanisms and that transmission acts as a filter to select for resistance with a lower cost or lower chance of reversion. Thus, when we observe resistance strains transmitting in the hospital, we propose that the resistant strains are less likely to revert to sensitivity, either because of compensatory mutations or the filtering mechanism.

To assess our hypothesis, we test the prediction that antibiotic resistance arising *de novo* will revert more readily than transmitted resistance. We identify a collection of daptomycin-resistant clinical isolates from a hospital system that experienced a rise in cases of daptomycin-resistant *E. faecium* infections (*4*). The isolates were drawn from two groups: those that were resistant at the time of the patient’s first positive blood culture (putatively transmitted strains) and isolates that were initially sensitive on the patient’s first blood culture but had resistant isolates following daptomycin treatment (*i.e.* potentially *de novo* resistance). We begin by characterizing the resistance phenotype and genetic pathways of resistance from these two groups. We then experimentally evolve these isolates in antibiotic-free medium, tracking both the phenotypic changes and genetic changes using whole genome sequencing. Our work suggests that antibiotic-resistant isolates that reverted rapidly do so because their resistance mutations also resulted in readily accessible mutational pathways for reversion to susceptibility. This shared evolutionary pathway may be an important process constraining the spread of high-level daptomycin resistance of *E. faecium*.

## 3 Materials and Methods

### 3.1 Selection of Clinical Isolates

Bacterial isolates were obtained from the University of Michigan Clinical Microbiology Laboratory and classified as daptomycin sensitive (MIC*≤* 4µg*/*mL) or resistant (MIC*>* 4µg*/*mL) using the Trek diagnostic system (ThermoFisher). Isolates were obtained from six patients with *de novo* resistant *E. faecium* bacteremias (hereafter DN), where the first isolate was daptomycin susceptible, but one or more subsequent isolates were identified as daptomycin resistant. Isolates were also obtained from six patients with a putatively transmitted resistant bacteria (hereafter PT), where a daptomycin-resistant, *E. faecium* bacteremia was identified on their first culture. For each DN patient, the earliest dated sensitive (DNS isolates) and resistant (DNR isolates) isolates were selected for further study. For each PT patient, the initial, daptomycin-resistant isolate (PTR isolates) was selected for further study. In total, 18 patient-derived isolates (6 PTR, 6 DNR, 6 DNS) were used. Isolates from patient PT6 were classified as sensitive by the clinical microbiology lab, but among the more daptomycin resistant internally tested isolates and therefore considered resistant for this study.

### 3.2 Laboratory evolution

A single colony from each blood culture sample was isolated on Brain Heart Infusion (BHI) agar (Becton, Dickinson and Company, 211065) and stored in BHI broth (Becton, Dickinson and Company, 237500) supplemented with 20% glycerol at -80*^◦^*C. To initiate experimental populations, a single clone from each isolate was streaked onto BHI agar and grown overnight at 36*^◦^*C. Thus, we expect no shared genetic variation between evolved replicate populations. Three replicate populations for each of the 18 clinical isolates were inoculated from the single colony. Cultures were grown in 10mL BHI broth in test tubes (18×150mm, Fisher 14-961-32), incubated at 36*^◦^*C. Daily transfers were performed by transferring 10 µL from the overnight culture into 10mL fresh BHI broth and were continued for 32 days. Transfers were interrupted for one day on Day 17 and re-started in 10mL of the Day 17 stock frozen in 20% glycerol stored at -80*^◦^*C. On the final day, three random clones were chosen from each population for subsequent genetic and phenotypic testing.

### 3.3 Population assessment of daptomycin resistance during experimental evolution

Evolving populations were assayed for susceptibility to daptomycin on days 8, 10, 17, 18, 25, and 32. A well mixed sample of each population after each growth cycle was serially diluted from 10^0^ to 10*^−^*^5^ in 10-fold steps with 0.85% saline. A 100 µL sample of each dilution was plated onto cation-adjusted Mueller-Hinton agar (BD Difco, catalog number. 212322) agar prepared with no daptomycin and 8 µg*/*mL daptomycin supplemented with 50 µg/mL Ca^2+^. Plates were incubated at 36 *^◦^*C for 24 hours. Colony forming units (CFUs) were counted at each concentration level.

### 3.4 Daptomycin minimum inhibitory concentration calculation

Daptomycin resistance was measured with a broth micro-dilution assay. Mueller-Hinton broth supplemented with 50 µg*/*mL Ca^2+^ and daptomycin concentrations ranging from 0.5 µg*/*mL to 16 µg*/*mL in serial two-fold increments. Each assay was performed in duplicate. Optical density readings were taken at 600 nm (OD_600_) at 21 hours (FLUOstar Omega, BMGTech) and a custom R script was used to fit a Hill function to calculate the minimum inhibitory concentration (MIC) as described previously (*19*). We defined the MIC as the concentration at which the OD is predicted to drop below the detection limit. Three reference clones, having low, medium, and high daptomycin resistance, were used as standard MICs to confirm internal consistency between blocks.

### 3.5 Statistical analysis of resistance phenotypes

A comparison of antibiotic resistance between groups was performed using linear mixed-effects models. The dependent variable was log_2_(MIC_evolved_ *−* MIC_anc_), where MIC_evolved_ is the calculated MIC of an evolved clone, and MIC_anc_ is the calculated MIC of the isolate from which it was experimentally evolved. Fixed effects included the evolved group (DNR, DNS, PTR) and initial MIC. Random effects allowed for variation due to replicated MIC measurements, with each clone nested within the replicated experimental population and each replicated experimental population nested within the patient. Models were fit by maximizing the log-likelihood using nlme package v3.1.141 (*20*) in R v3.5.1.

### 3.6 Fitness estimation in laboratory media

Fitness estimates were determined by analyzing growth curves of individual clones over 20 hours. Frozen stocks of clones were inoculated into BHI liquid media, the same media used in the evolution experiment, and grown overnight. Clones were then diluted 1:1000 in BHI media and 100 µL of each sample was aliquoted into 96 well flat-bottom plates (Celltreat). OD_600_ measurements were taken using an automatic plate reader (FLUOstar Omega, BMGTech) every 15 minutes for 20 hours. Plates were shaken at 300 rpm for 1 minute prior to each reading.

The OD_600_ measurements were then fitted to a five parameter logistic growth model as described by (*21*). The growth model is as follows:

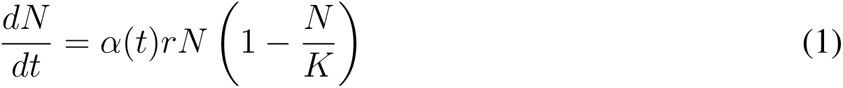

where N is the density of cell growth as measured by OD_600_ measurements, *α*(*t*) is a logistic function of two parameters representing the proportion of cells actively dividing, r is the maximum growth rate, and K is the carrying capacity. Relative fitness was estimated assuming competition for a single limiting resource with no interaction between strains (*22*). The ances-tral clone from patient PT4 was used as an arbitrary reference for all relative-fitness calculations. Blocks were structured such that initial and experimentally evolved clones from one DN patient and one PTR patient were measured on the same plate. Each clone was independently grown on three different days. Five wells from each plate contained only growth media, which served as blanks.

Rather than fitting the model with least squares as done previously (*22*), we use the POMP package (*23*) to better account for sources of measurement error across the growth cycle. Specifically, we used log-normal errors to account for multiplicative errors arising from cell clumping occurring at the end of the growth cycle. For each replicate, growth model parameters were initialized using a range of starting values. Parameter values were then optimized using the Nelder-Mead algorithm from the optim function of the stats package to identify the maximum likelihood estimate of each parameter. More details can be found in the supplement.

### 3.7 Sequencing Library preparation

Each of the 18 clones used to found experimentally evolved populations were sequenced using both long read (MinIon MIN-101B, Oxford Nanopore) and short read (NovaSeq 6000, Illumina) technology. Nanopore sequencing libraries were prepared using the SQK-LSK109 kit according to manufacturer protocols. Long read samples were multiplexed in two libraries containing 4 and 14 samples and barcoded with the EXP-NBD104 Kit. The first library of 4 samples was run for 6 hours using a MIN-101B sequencer on a MinION R9.4.1 flow cell. The flow cell was then washed using the EXP-WSH003 flowcell wash kit. The second library of 14 samples was then loaded and allowed to run for 24 hours. Illumina sequencing libraries were prepared using an Illumina TruSeq DNA LT kit according to manufacturer protocols. The short and long read sequencing data is available on NCBI Bioproject PRJNA975905 with labels as indicated in Table 1.

**Table 1:** Associated metadata of the study isolates and their respective phenotype, MLST, study grouping, NCBI accession, and isolate number.

| Patient | Phenotype | Group ID | MLST | Isolate Number | Accession | Clinical MIC ($\mu\text{g/mL}$) |
| --- | --- | --- | --- | --- | --- | --- |
| DN1 | Sensitive | DNS | 412 | BL00201-1 | SAMN35347354 | 2 |
| DN2 | Sensitive | DNS | 17 | BL00216-1 | SAMN35347358 | 4 |
| DN3 | Sensitive | DNS | 18 | BL00239-1 | SAMN35347362 | 4 |
| DN4 | Sensitive | DNS | 584 | BL00242-1 | SAMN35347366 | 4 |
| DN5 | Sensitive | DNS | 17 | BL00244-1 | SAMN35347370 | 2 |
| DN6 | Sensitive | DNS | 18 | BL00247-1 | SAMN35347374 | 4 |
| DN1 | Resistant | DNR | 412 | BL00211-1 | SAMN35347378 | 8 |
| DN2 | Resistant | DNR | 17 | BL00223-1 | SAMN35347382 | 32 |
| DN3 | Resistant | DNR | 18 | BL00241-1 | SAMN35347386 | 8 |
| DN4 | Resistant | DNR | 584 | BL00243-1 | SAMN35347390 | 8 |
| DN5 | Resistant | DNR | 17 | BL00246-1 | SAMN35347394 | 16 |
| DN6 | Resistant | DNR | 18 | BL00250-1 | SAMN35347398 | 16 |
| PT1 | Resistant | PTR | 1471 | BL00192-1 | SAMN35347330 | 16 |
| PT2 | Resistant | PTR | - | BL00194-1 | SAMN35347334 | 8 |
| PT3 | Resistant | PTR | 1471 | BL00196-1 | SAMN35347338 | 8 |
| PT4 | Resistant | PTR | 664 | BL00198-1 | SAMN35347342 | 8 |
| PT5 | Resistant | PTR | 412 | BL00184-1 | SAMN35347346 | 8 |
| PT6 | Resistant | PTR | 664 | BL00777-1 | SAMN35347350 | 4 |

### 3.8 Processing and filtering of sequence data

For each sample, Illumina sequencing adapters were trimmed and reads with PHRED score *<* 25 or were unpaired removed using Trimmomatic v0.38 (*24*). Quality checks of the original and trimmed data were performed using FastQC v0.11.8 (*25*). For Nanopore sequencing data, Albacore v2.3.4 (https://community.nanoporetech.com/downloads) was used to call bases and de-multiplex. Nanopore sequencing adapters were trimmed using Porechop (https://github.com/rrwick/Porechop/), and reads with lengths less than 500 bases were removed using NanoFilt v.2.7.1 (*26*).

### 3.9 Genome assembly and annotation

Genome assembly was performed using the hybrid-assembler Unicycler v0.4.4 with normal bridge settings (*27*). Multi-locus sequencing types (MLST) were determined using mlst software (https://github.com/tseemann/mlst) against the pubMLST database (*28*). Prokka v1.14.6 was used to annotate the genome assemblies (*29*).

### 3.10 Variant calling

Short read sequencing reads were mapped to the assembly of their respective ancestor using bwa-mem v0.7.17 (*30*). Variant calling was performed using the Genome Analysis Toolkit (GATK4) v4.1.2 (*31*). Variants occurring on plasmids were removed. Insertion sequence elements were identified using panISa (*32*). Variants were then annotated using snpEff v4.3t (*33*) using the annotation of the ancestral clone generated by Prokka (*29*). All variants were visually inspected using the Integrative Genomics Viewer (IGV) (*34*). For the croRS mutation occurring between DNS2 and DNR2, a structural variant was found using GATK. This mutation was later identified as an inversion using long read sequencing (*35*). Lollipop plots used to display variants were generated using trackViewer (*36*).

### 3.11 Mutations in candidate daptomycin resistance genes

To identify mutations in genes previously associated with daptomycin resistance, short-read sequences from each isolate were mapped to the published sequence of reference strain DO (CP003583) (*37*) and called using the protocol above. Mutations were considered candidate daptomycin resistance gene mutations if they were in genes previously associated with daptomycin resistance (*38*). Additionally, variants were removed if present in multiple MLSTs as resistance was not associated with MLST in prior analysis (*35*). Genes annotated as hypothetical or phage-associated were also removed from the list of considered genes due to the high level of variability and low quality of evidence. The mutations are listed in Table 2.

**Table 2:**
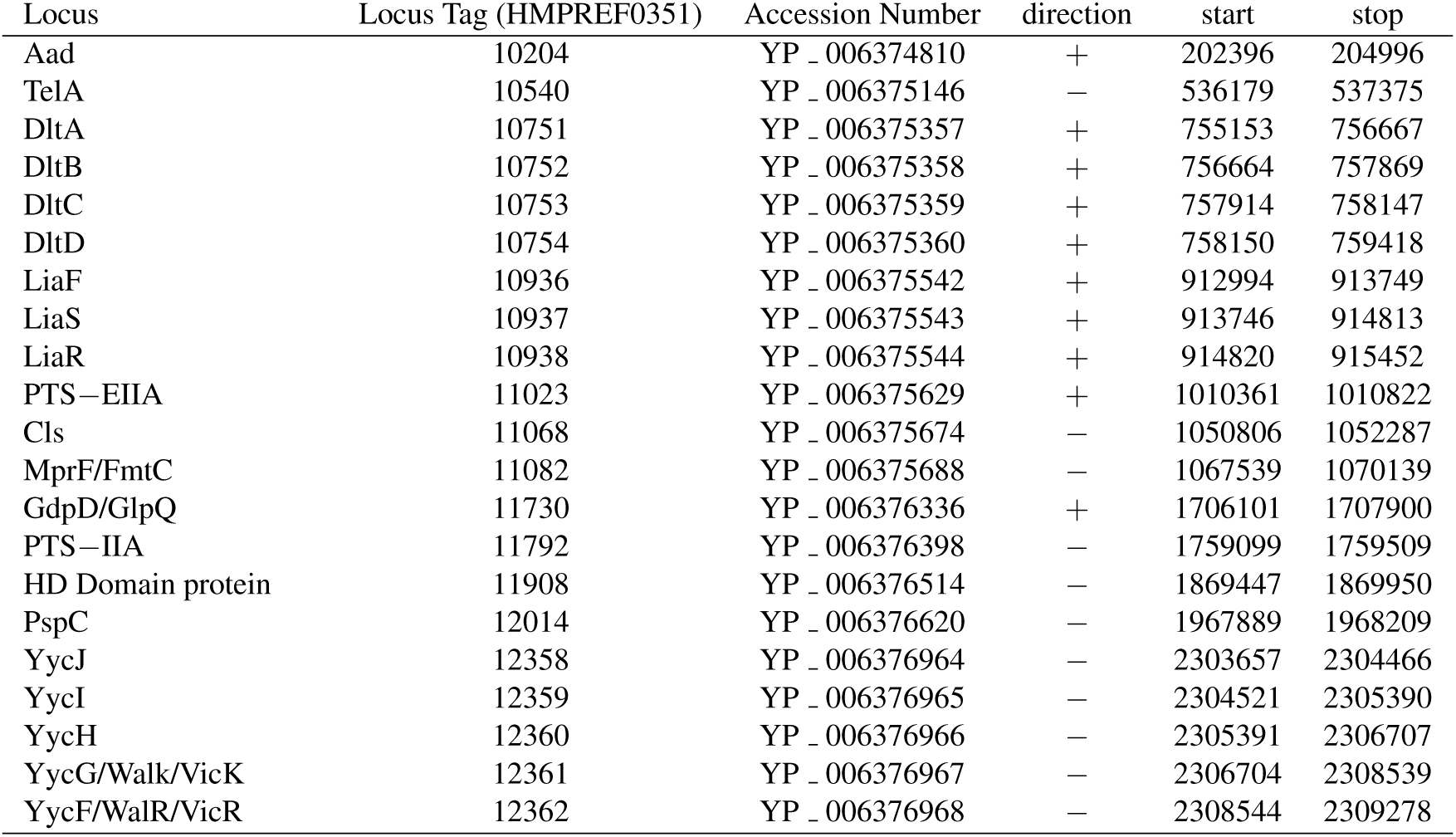
Candidate daptomycin resistance genes as reported in Diaz et al. (***38***) **with mutations occurring across multiple MLSTs removed** (***35***).

### 3.12 Phylogenetic tree estimation of initial isolates

For the 18 initial clones, a core genome was created using snippy-core (https://github.com/tseemann/snippy) and then converted to a single nucleotide polymorphism (SNP) distance matrix using snp-dists 0.7.0 (https://github.com/tseemann/snp-dists). A neighbor-joining tree (*39*) was created in R v4.0.2 using the APE (*40*) package.

### 3.13 Data and code availability

The data and code used in this study are available at https://osf.io/8hsn2/.

## 4 Results

### 4.1 Identification of strains and isolates for the evolution experiments

Patients were classified as either *de novo* resistant (DN) or putatively transmitted (PT). DN patients had an initial blood culture grow daptomycin susceptible *E. faecium*. After daptomycin treatment, a subsequent blood culture grew daptomycin resistant *E. faecium*. PT patients had an initial positive blood culture grow a daptomycin resistant *E. faecium*. The isolates derived from these patients were labeled according to the patient classifier (DN or PT), the resistant status, and a unique number. For example, DNS1 and DNR1 are the sensitive and resistant isolates from patient DN1. The DNR isolates, DNS isolates, and putatively transmitted resistant isolate (PTR) were further studied using experimental evolution in antibiotic free conditions (Figure 1). The patient, clones, MLST, and clinical MIC for each initial isolate are listed in Table 1.

**Figure 1:**
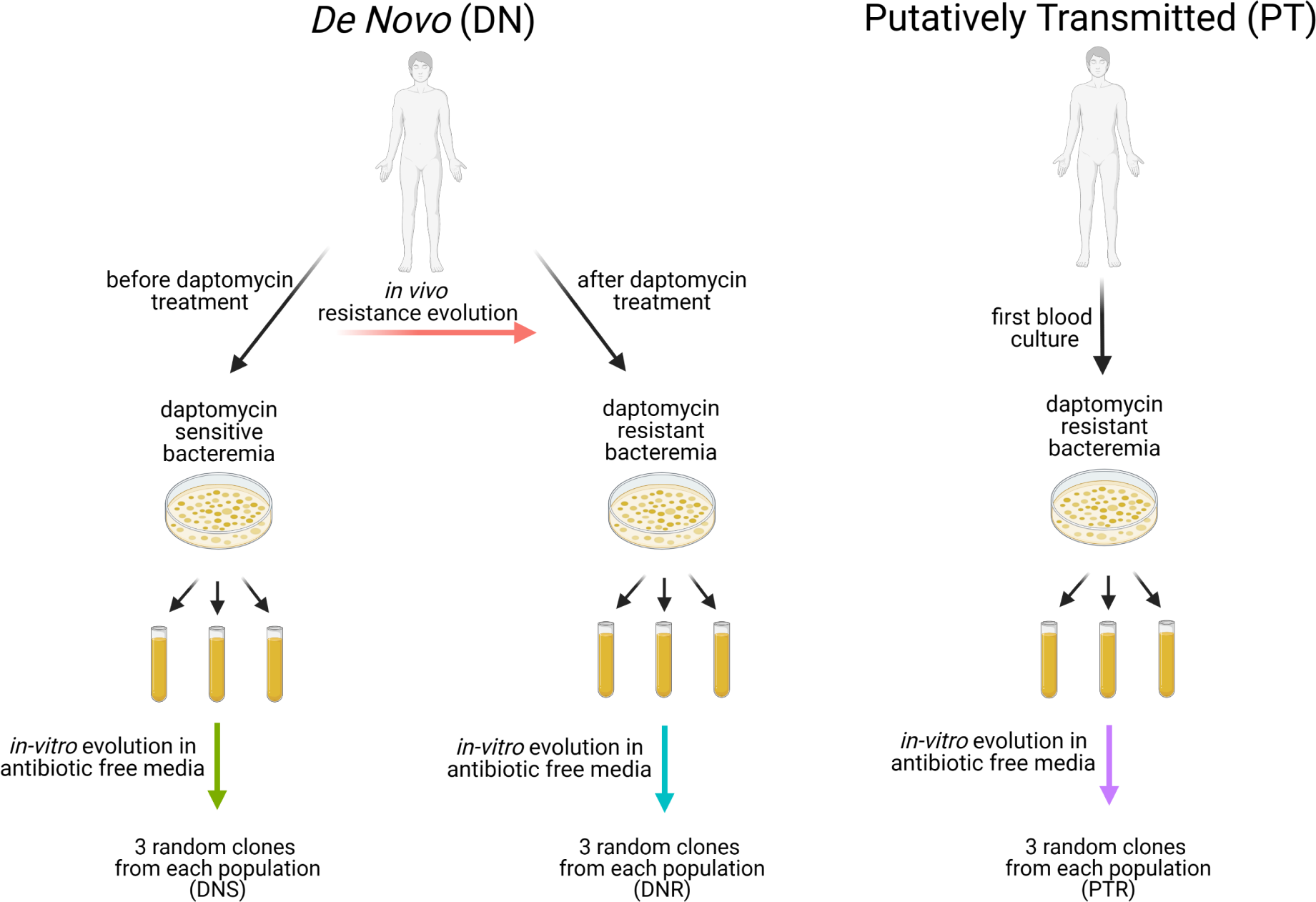
Experimental Design. Isolates were obtained from patients with one of two types of VRE bacteremia: de novo resistant patients that converted from having daptomycin sensitive to daptomycin resistant VRE bacteremia or putatively transmitted resistance patients with daptomycin resistant on their initial culture. Three independent populations were evolved from the initial daptomycin sensitive strains, the converted daptomycin resistant strain, and the daptomycin strains from patients resistant on arrival for 32 days in BHI media. Figure was created using Biorender.com.

### 4.2 Resistance levels of *de novo* resistant and transmitted strains

The mean calculated MIC of the PT strains was 4.65 µg/ml (range 1.46 to 14.7), while the calculated MIC of the DN resistant strains was 10.3 µg/ml (range 1.22 to 45.3) (Figure 9). Given the small sample size, this difference is not unexpected (p=0.31, Mann-Whitney U). Reassuringly, the resistant isolate of the DN pairs was more resistant than their corresponding sensitive relative (p=0.031, Wilcoxon sign rank test). As a group, sensitive isolates from the DN pairs had MICs significantly lower than resistant isolates in both DN and PT patients (p=0.002, Mann-Whitney U). As in prior studies, there were differences between MICs determined by the clinical microbiology laboratory and those determined using laboratory protocols described above (*4*). Some strains classified as daptomycin resistant by the clinical lab were experimentally assayed MICs below the CLSI breakpoint of resistance (MIC *>* 4µg/ml), a reflection of the difference in assays and the continuous nature of our MIC measurement.

### 4.3 Experimental evolution in antibiotic-free media reveals different patterns of adaptation

Daptomycin resistance was assessed periodically by spreading samples from each experimentally evolving population onto agar plates with and without daptomycin (Figure 2). In five of the six populations founded with the resistant clone from an DN patient, there was a consistent decrease in the ratio of resistant colonies among all three populations by the end of the evolution experiment(Figure 2). In contrast, a decline across all three populations was observed from only one PT ancestor (PT6). We also note that none of the populations from PTR2 showed any growth on daptomycin plates. This lack of growth for a resistant strain was not entirely unexpected, given differences in testing methodology and the high variability in testing daptomycin MICs (*41*).

**Figure 2:**
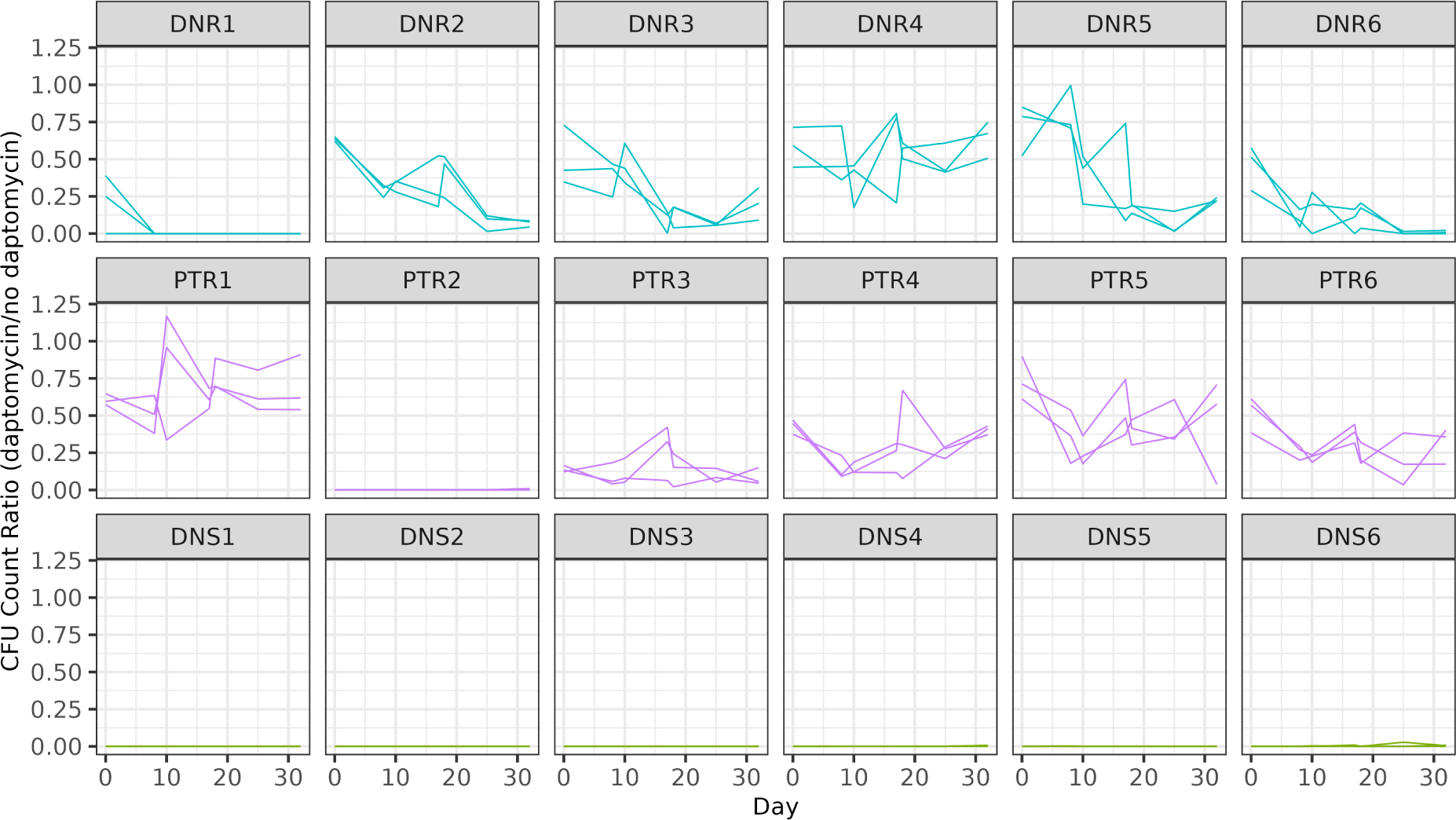
Population level of resistance during experimental evolution. Resistance is measured as the ratio of colonies growing on plates supplemented with daptomycin at 8 µg*/*mL to colonies on plates without antibiotic for each of three independent experimental populations evolved from each patient-derived isolate. Populations were tested on days 0, 8, 10, 17, 18, 25, and 32.

We next selected three random clones from the final time point of each experimental population and measured their daptomycin MIC. We assess the hypothesis that resistance that recently evolved *de novo* within-host would revert more than the resistance that is transmitted between hosts. We compared the relative changes in MICs in the DNR and the PTR using a linear mixed model (Table 3). The patient group was not a significant predictor of the relative change in MIC (Model 1, p=0.26). However, the initial MIC of the founding clone was a highly significant predictor of relative change in MIC (Model 2, p*<*0.001).

**Table 3:** Maximum likelihood fit of three mixed models, as described in the text, to estimate the effect of group (*i.e.* being founded with a PT or DN clone), and the MIC of that founding clone, on the reduction in MIC after 32 days of evolution in antibiotic-free conditions. difference (individual) as output, does NOT include DNS group

|  | Value | Std.Error | DF | t-value | p-value |
| --- | --- | --- | --- | --- | --- |
| **MODEL 1: $\log_2(\text{final MIC} - \text{initial MIC}) \sim \text{Group}$** |  |  |  |  |  |
| Intercept | -0.6999106 | 0.5890485 | 108 | -1.19 | 0.24 |
| Group | -1.0040332 | 0.8330403 | 10 | -1.2052635 | 0.2558477 |
| **MODEL 2: $\log_2(\text{final MIC} - \text{initial MIC}) \sim \text{Group} + \log_2(\text{initial MIC})$** |  |  |  |  |  |
| Intercept | 1.2206256 | 0.3132934 | 107 | 3.90 | 0.0002 |
| Group | -0.2835307 | 0.3981069 | 10 | -0.71 | 0.49 |
| $\log_2(\text{initial MIC})$ | -1.0170222 | 0.0755229 | 107 | -13.47 | <0.001 |
| **MODEL 3: $\text{MIC}_{\log 2} \sim \text{Group} + \text{InitMIC}_{\log 2} + \text{Time}$** |  |  |  |  |  |
| (Intercept) | 1.8352468 | 0.2789396 | 143 | 6.58 | <0.001 |
| GroupDNR | -0.2057706 | 0.2972867 | 10 | -0.69 | 0.50 |
| InitMIC<sub>log2</sub> | 0.2275255 | 0.0634888 | 143 | 3.58 | 0.0004 |
| Time | -0.0400642 | 0.0063254 | 107 | -6.33 | <0.001 |

While we did not find a significant difference in the initial MICs between DNR and PTR isolates, isolates with higher MIC tended to belong to DNR and show greater absolute declines in MIC. Populations founded with resistant isolates from four of the DN patients (DN1, DN3, DN5, DN6) and one PT patient (PT5) had a decline of more than 2 µg/ml, when averaged across the three replicate populations. In contrast, populations evolved from three PT isolates (PT1, PT2, PT4) had no significant change (Figure 9). Four of the six resistant clones from DN patients (DN2, DN4, DN5, and DN6) had at least one of their evolved populations return to MIC near their sensitive pair. Finally, all three clones from all three populations evolved from the resistant isolates from DN1 were even more sensitive than their sensitive paired ancestor.

### 4.4 Growth curves and single resource competition model

We next estimated the *in vitro* fitness of isolates from growth curves (*22*). For each DN patient, we measured the fitness of the initial DNS, initial DNR, and 3 clones from each of the 3 evolved populations (figure 3). For each PT patient, we measured the fitness of the initial PTR and 3 clones from each of the 3 evolved populations. We find no significant difference between the initial fitness of DNS and DNR isolates (Table 4, p=0.0636, Model 1) or DNR and PTR isolates (p=0.697, model 2). Laboratory evolved clones had greater fitness than their ancestors (p*<*0.01, model 3). Fitness changes were influenced by starting MIC (model 4, p=0.0002) and starting fitness (p=0.0003, model 5). While the trend of increased fitness after evolution was consistent across strains, we again note that a substantial decrease in fitness was found in PTR4 after evolution. Repeating the analysis excluding PTR4 does not significantly change model results.

**Figure 3:**
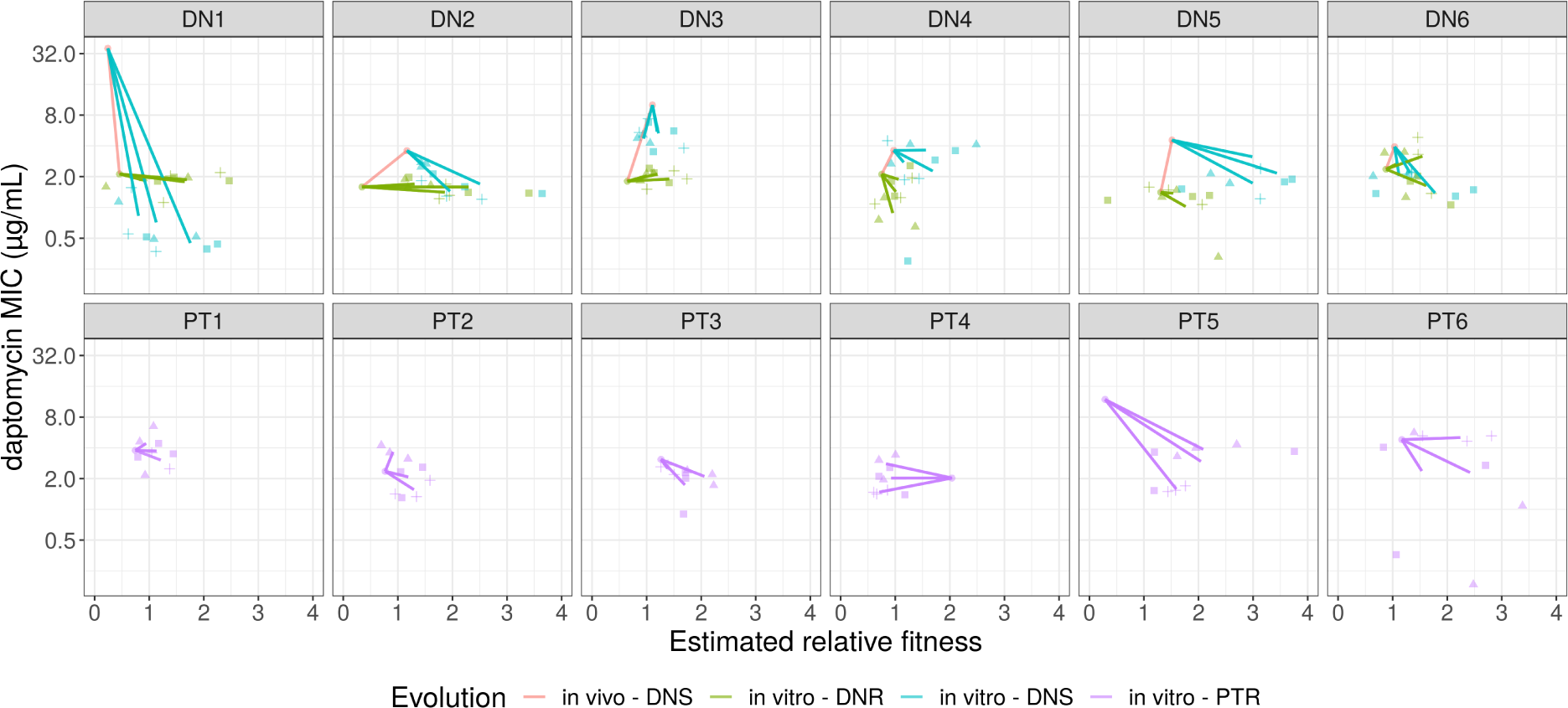
MIC and fitness relationship. The relationship between the mathematically modelled fitness of a clone (see methods) and its MIC. For each PTR, DNR, and DNS isolate, the fitness and MIC of the initial clone and three clones from three experimental evolved replicate populations were estimated. For DN patients, the change from the initial DNS to the DNR clone or the difference between the DNS and the DNR clones was also estimated. Each color represents a group, with the red line representing *in vivo* changes between initial DNS and DNR isolates. The remaining lines represent the average changes from the initial clone to the average of the three clones from each replicate population. Colored shapes show the individual measurements of each clone from each replicate population.

**Table 4:**
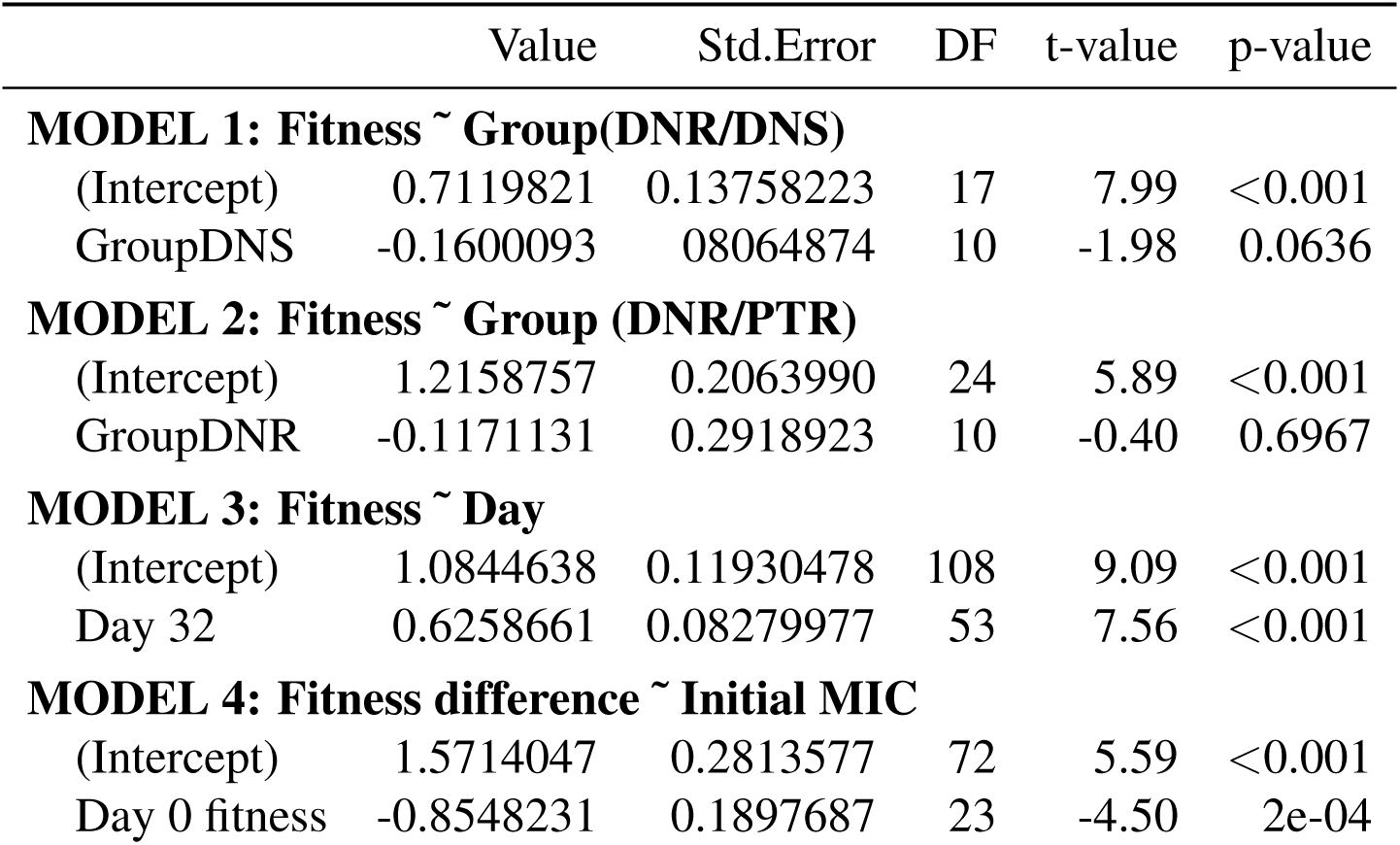
Maximum likelihood fit of mixed models to estimate the effect of group (*i.e.* being founded with a PT or DN clone), fitness gain during adaptation, and starting MIC on the fitness of strains after 32 days of evolution in antibiotic-free conditions.

|  | Value | Std.Error | DF | t-value | p-value |
| --- | --- | --- | --- | --- | --- |
| **MODEL 1: Fitness ~ Group(DNR/DNS)** |  |  |  |  |  |
| (Intercept) | 0.7119821 | 0.13758223 | 17 | 7.99 | <0.001 |
| GroupDNS | -0.1600093 | 0.08064874 | 10 | -1.98 | 0.0636 |
| **MODEL 2: Fitness ~ Group (DNR/PTR)** |  |  |  |  |  |
| (Intercept) | 1.2158757 | 0.2063990 | 24 | 5.89 | <0.001 |
| GroupDNR | -0.1171131 | 0.2918923 | 10 | -0.40 | 0.6967 |
| **MODEL 3: Fitness ~ Day** |  |  |  |  |  |
| (Intercept) | 1.0844638 | 0.11930478 | 108 | 9.09 | <0.001 |
| Day 32 | 0.6258661 | 0.08279977 | 53 | 7.56 | <0.001 |
| **MODEL 4: Fitness difference ~ Initial MIC** |  |  |  |  |  |
| (Intercept) | 1.5714047 | 0.2813577 | 72 | 5.59 | <0.001 |
| Day 0 fitness | -0.8548231 | 0.1897687 | 23 | -4.50 | 2e-04 |

### 4.5 Genetic backgrounds of clinical isolates

Bacterial chromosomes were fully closed and circularized for 13 of the 18 clinical isolates, while the remaining 5 isolates had at least one contig *>* 2.7Mb. These isolates were genetically diverse, with multiple MLSTs represented (Table 1). All strains were separated by 40 or more single nucleotide polymorphisms (SNP), except strains PT1 and PT3 which were separated by 11 SNPs. Strains DNR1 and DNS1 (sensitive-resistant pairs) were separated from strain PT5 by 45 and 53 SNPs, respectively.

Mutations in genes previously associated with daptomycin resistance (*38*) were found in most, but not all clinically resistant isolates (Table 2). Resistant strains from patient DN4 and DN6 had mutations in *clsA*, strains from patient DN5 had a *glpQ* mutation, and strains from patient DN1 had *liaS*, *HD Domain*, and *yycG* mutations (Figure 4). PTR isolates do not have a sensitive ancestor for genetic comparison. We therefore identify differences in daptomycin resistance associated genes relative to the sensitive *E. faecium* reference genome DO. Three of six PTR isolates showed mutations in either *clsA* or *liaS*, which are most strongly associated with daptomycin resistance (Figure 5).

**Figure 4:**
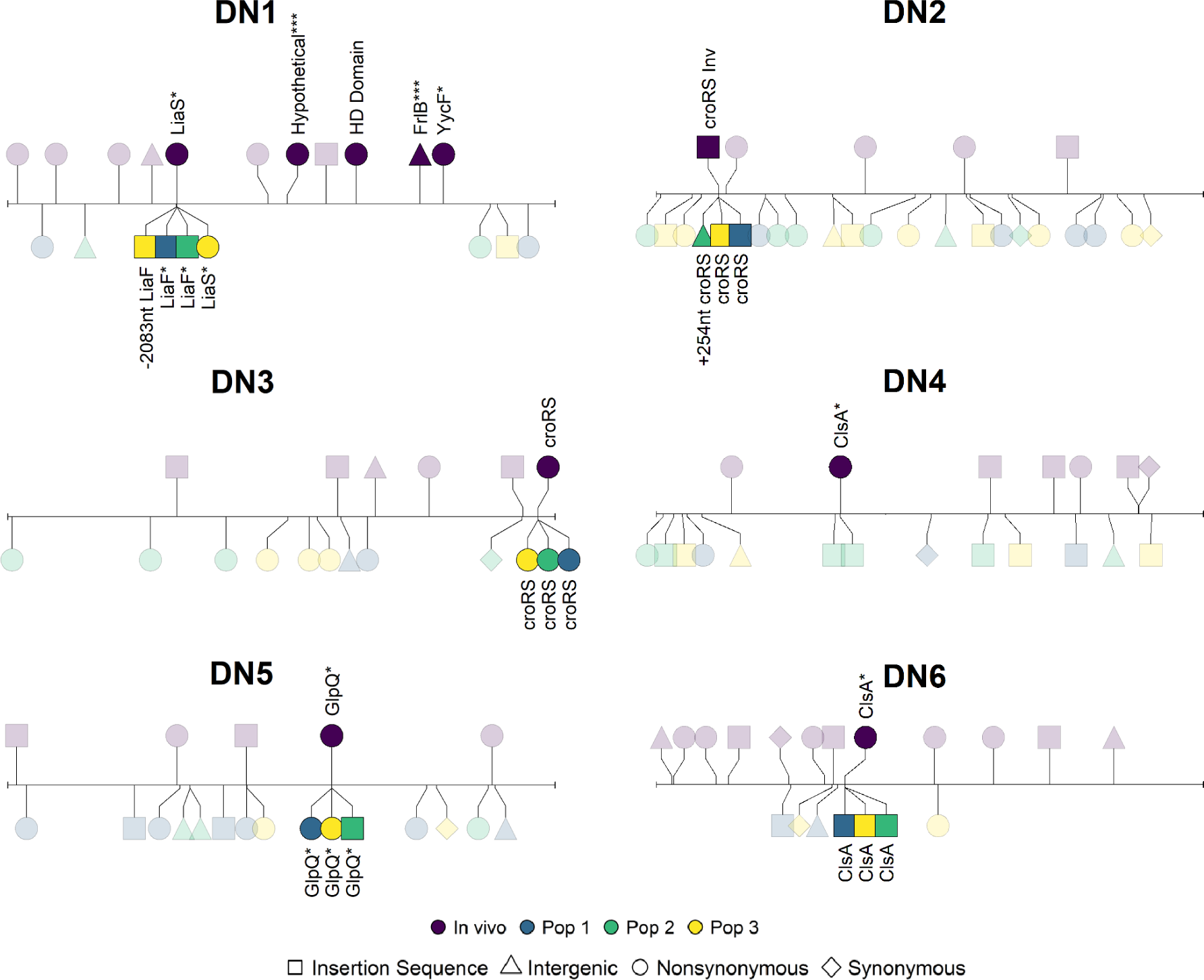
**Mutations in laboratory evolved populations from patients with *de novo* resistance**. Lollipop plot showing mutations occurring during *in vivo* and *in vitro* evolution. The line represents the chromosome of the sensitive isolate and each lollipop represents a mutation. Differences between the resistant and sensitive isolate (*in vivo) evolution* are above the chromosome, while mutations seen during experimental evolution are below. The color indicated the experimental replicate and the shape indicated the mutation class. Genes or operons with mutations in multiple experimental populations or when the gene has been associated with daptomycin resistance are labeled. Mutations labeled with one asterisk are mutations that have been previously associated with daptomycin resistance. Mutations labeled with three asterisks in DN1 appeared in the initial resistant isolate and only one of the evolved resistant populations. See text for further explanation.

**Figure 5:**
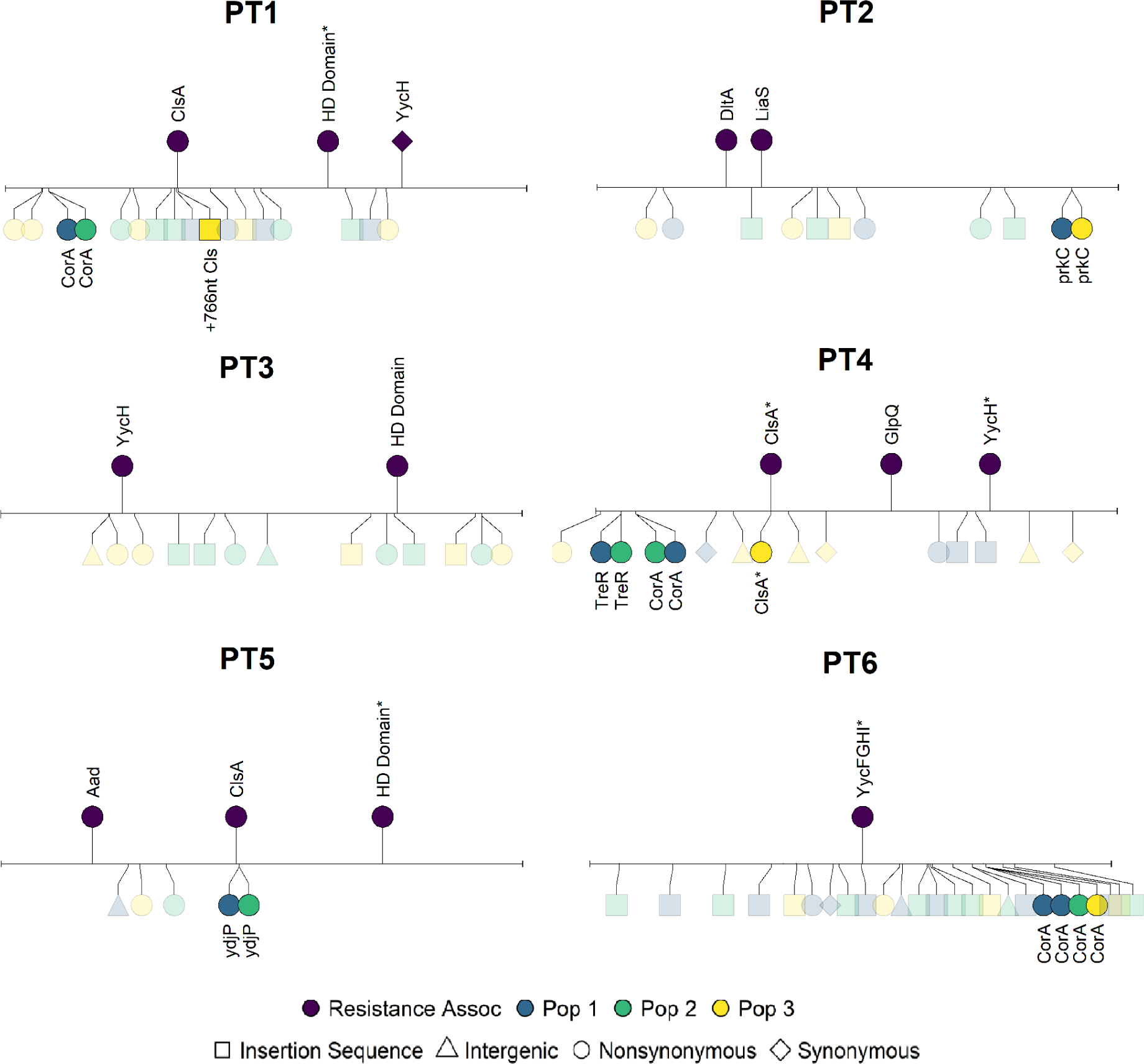
Mutations in laboratory evolved populations from transmitted resistance patients. Lollipop plots showing genetic changes occurring in isolates from PT patients. Lollipops on the top half of the plot show mutations in genes associated with daptomycin resistance mapped against the daptomycin sensitive reference genome DO. Gene names with asterisks have multiple mutations within the gene. Lollipops on the bottom half of the plot show genetic differences in one clone selected from each of the three experimentally evolved populations.

### 4.6 Genetic evolution in experimental populations

We identified 0-10 genetic changes in the laboratory evolved clones relative to the founding clone for each population, with a mean of 3.8 (IQR 2.5-5), though this number differed between the *de novo* sensitive, *de novo* resistant, and the transmitted groups (3.1 vs 3.7 vs 4.6 respectively).

Parallel evolution consisting of mutations in the same gene or operon across replicate laboratory evolved populations from the same founding genotype was observed in some cases. This parallelism was more common in the DNR isolates than in the PTR isolates (5 out of 6 DNR versus 1 out of 6 PTR). Notably, the site of parallel evolution varied across founding genotypes and in all of the DNR founded populations. Mutations in these populations were associated with genes or operons previously suggested to be resistant isolates (*35*), whereas the one case of parallel evolution from a PTR isolate was not.

We found that for DNR1, the genotype of the initial resistant clone used to found the replicate populations differed between populations. This was identified when two mutations in DNR1 were not found in population 1. Resequencing of the DNR1 population revealed these two mutations were polymorphic in what had been assumed to be a clonal population. Thus, we suspect the initial resistant isolate used to found populations 2 and 3 did not carry these two mutations not found in population 1. These mutations are marked in Figure 4 and have not been linked to daptomycin resistance.

Among PTR populations, one replicate population from PTR4 showed a second mutation within *ClsA*. In addition, one replicate population from PT1 had an insertion sequence arise near *ClsA*. No other experimentally evolved populations from transmitted resistant isolates had mutations in resistance associated genes.

Parallel mutations in the magnesium transporter *corA* were commonly seen in populations founded by sensitive and resistant isolates suggesting adaptations to laboratory conditions independent of daptomycin resistance.

## 5 Discussion

Under strong antibiotic pressure, most bacteria can evolve resistance to antibiotics. Yet, most of these adaptations are evolutionary dead-ends once antibiotic pressure is removed or the host clears the organism (*7*). Antibiotic resistance arising within hosts and spreading between hosts is rarer but far more concerning. This study examined isolates that recently evolved *de novo* resistance within host or transmitted between hosts. We find significant differences in their phenotypic robustness in antibiotic free conditions that likely constrain the spread of daptomycin resistant *E. faecium*.

The experimental evolution of these two groups of resistant bacteria in antibiotic free laboratory conditions identified three consistent and related patterns. First, the more resistant the founding isolate, the more the populations evolved towards sensitivity during the experimental period. Second, the three independent replicate populations from isolates in the *de novo* group often showed parallel molecular evolution with mutations occurring in the same gene or operon. Third, genes involved in the parallel evolution were unique to each resistant isolate and often occurred as a second mutation in the same gene or operon associated with resistance that evolved within the patient. These parallel mutations were never precisely the same across replicate populations and no precise genetic reversions were observed. The parallel mutations were often highly disruptive, such as the insertion of IS elements, frameshifts, and nonsense mutations. In contrast, we did not observe patterns of parallel evolution in resistance associated genes among putatively transmitted isolates. Together, these findings support our hypothesis that transmitted resistance strains are less likely to revert.

Previous research has focused on *in vitro* fitness costs as a limit to resistance transmission and spread (*42,43*). Our results suggest that the spread of daptomycin resistance *E. faecium* may also be slowed due to the accessibility of evolutionary pathways. The evolutionary pathways of resistant isolates once antibiotics are removed appear influenced by the initial resistance mutations. Indeed, unique second mutations occurred in or around a resistance mutation among all replicate populations in five of six *de novo* isolates, resulting in increased fitness in antibiotic-free environments and decreased resistance levels. Together, this suggests that the reversion of resistance phenotype among DNR may have been facilitated by a large mutational target not present among transmitted isolates. This pattern held across different genes, including enzymes (*clsA* and *glpQ*) and regulatory elements (*liaFSR* and *croRS*). In contrast, the evolutionary trajectories of transmitted resistant strains were less predictable, with no clear patterns of parallel molecular evolution, smaller decreases in resistance, but similar increases in fitness. Further study of this mechanism at the hospital level is warranted.

This study also demonstrates the utility of experimental evolution for understanding the evolutionary potential of antibiotic-resistant bacteria. Using controlled, replicated evolution in identical environments with defined starting genotypes, we can assess how commonly a particular evolutionary pathway is accessed. While the experimental environment differs from the clinical environment, we find a similar evolutionary pattern *in vivo* in patient DN1. A blood culture obtained after daptomycin was stopped showed a second site LiaS E192* nonsense mutation (*35*), which is consistent with experimental results showing disruptive mutations in LiaF or LiaS in all three replicate *in vitro* populations.

This study has several limitations. A relatively small number of isolates were used; thus, not all genetic backgrounds and resistance pathways were represented. It is also not possible to definitively say that resistance was *de novo* or transmitted, as the apparent *de novo* resistance could have resulted from two independent transmission events from closely related donors. Similarly, the incomplete records from these patients mean it is possible that putative transmitted resistance had daptomycin exposure. Further, the experimental environment differs in many important ways from the typical environment experienced *in vivo* by *E. faecium*. The calculated fitness is, therefore, an inexact measure of competitive fitness even in the laboratory environment. Despite these limitations, all of which would tend to obscure a pattern of difference, we nonetheless observe these convergent evolutionary patterns.

The rise of antibiotic resistance remains a pressing public health threat. Understanding the complex evolutionary dynamics leading to widespread resistance is a high priority. This study identifies the phenotypic reversion of resistance through a common evolutionary pathway as a potentially important process in limiting the spread of daptomycin resistance in *E. faecium*. The results presented here emphasize that the long-term success of a resistant isolate may depend on more than the fitness cost of resistance but also on the resulting evolutionary potential of the isolates in the absence of the antibiotic.

## 6 Acknowledgements

This research was supported by funding to AFR from Eberly College of Science and Huck Institutes of the Life Sciences, Pennsylvania State University and to RJW from the National Institutes of Health (R01AI143852 and K08 AI119182). The funders had no role in study design, data collection and analysis, decision to publish, or preparation of the manuscript.

## 7 Supplemental Data and Figures

**Figure 6:**
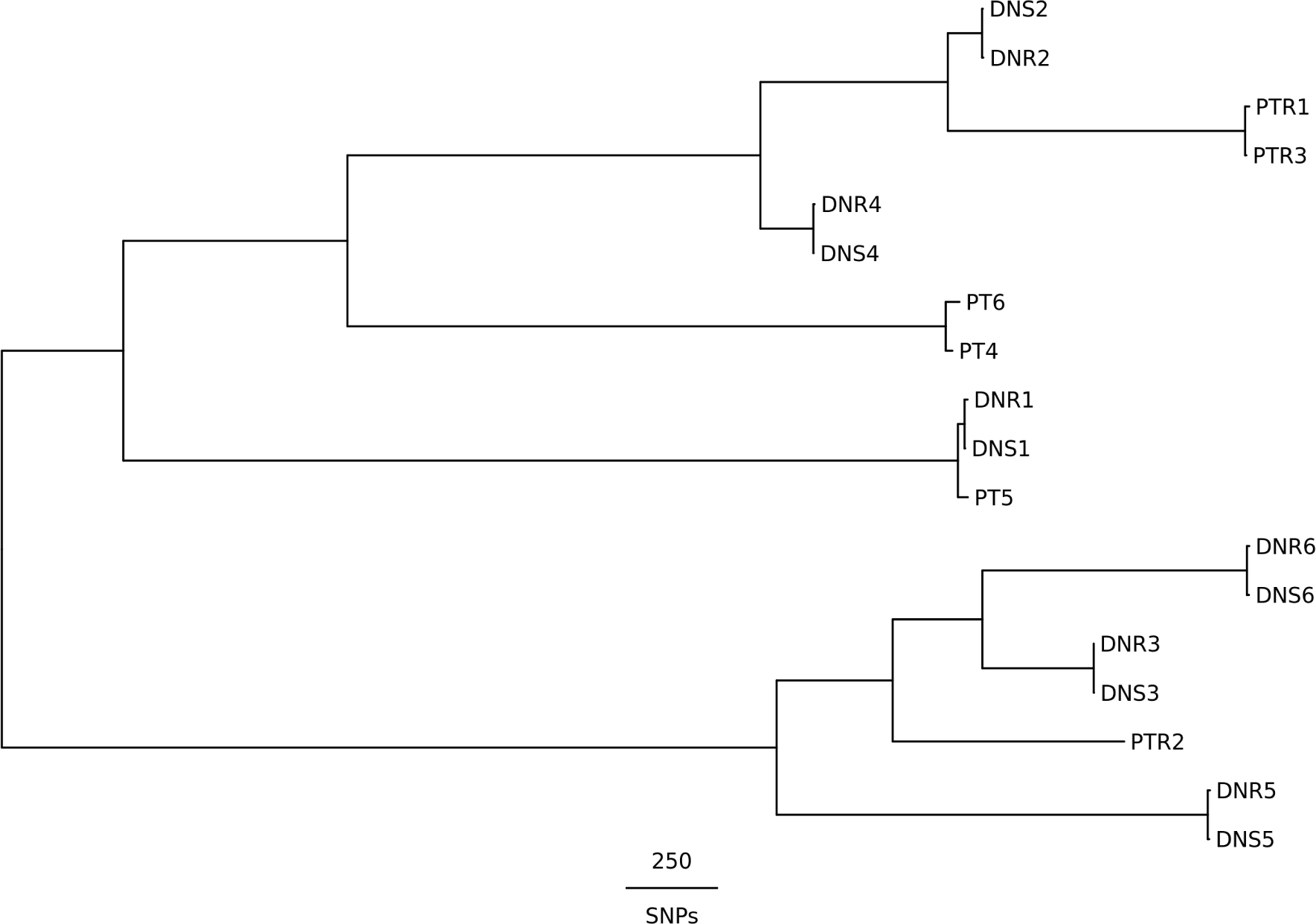
Phylogenetic tree of the 18 initial, unevolved isolates. A neighbor-joining tree of the initial 18 founding isolates from the SNP distance matrix. The SNP distance matrix was derived from a core genome alignment of all 18 isolates.

### Competition Model

We first fit the single species logistic growth model with a lag phase (*21*), where 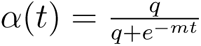. Parameters were estimated on each of the three clones taken from the 18 experimental population (6 DNR, 6 DNS, 6 PTR) across three biological replicates. Models were fit using POMP (*23*) with the Nelder-Mead optimization using the optim package. Models were fit assuming log-normal errors.

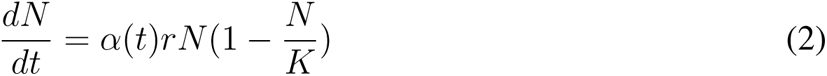

The parameter estimates taken from the single species model are then used to estimate the relative fitness values using the single resource competition model derived by Ram et al 2019 (*22*). Briefly summarizing, let R be the density of the limiting resource and N be density of cell populations. Cell growth is assumed to be proportional to R*·*N, resource is taken up by cells at rate h, and converted to cell mass at *ɛ*

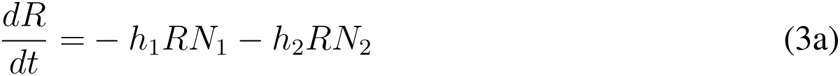

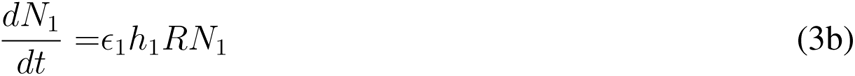

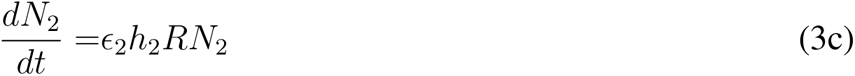

By conservation of mass:

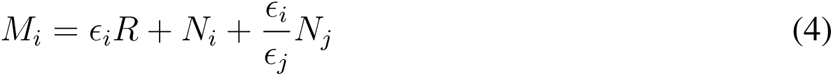

Since 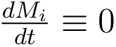 and *M_i_* is constant

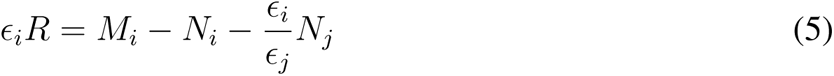

Substituting 5 into 3b and 3c, we get

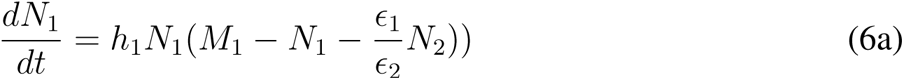

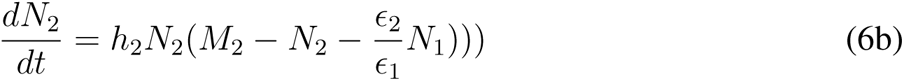

Let 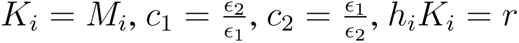

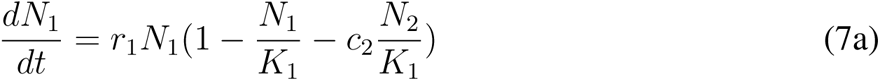

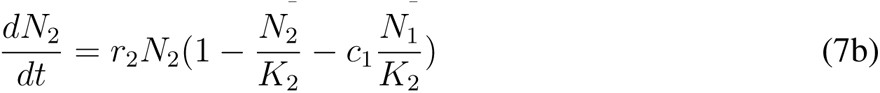

From the single species logistic growth:

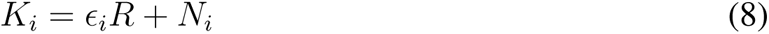

Ram et al assumes that *c*_1_ = *c*_2_ = 1 is an appropriate approximation based on empirical data. This assumption causes bacterial populations to decline if K is significantly different between the two species. This also implies that differences in R are driving the difference between *K*_1_ and *K*_2_, rather than a biological process

Rather we assume that the resources between the two single species systems are equal

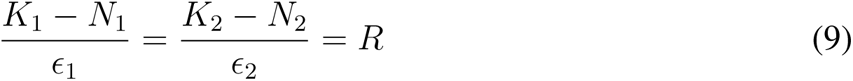

We can simplify this expression by assuming

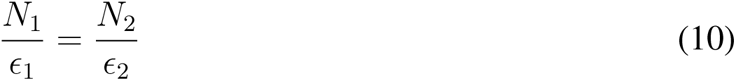

This provides the approximation

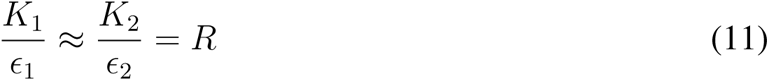

And we can then calculate *c*_1_ and *c*_2_ in terms of *K*_1_ and *K*_2_

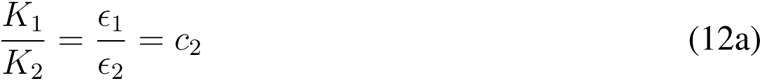

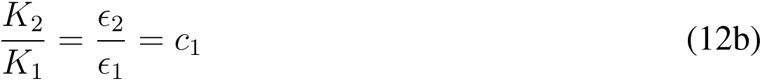

We then add the lag phase term to the equation to give us

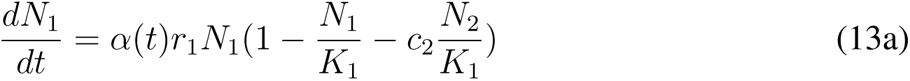

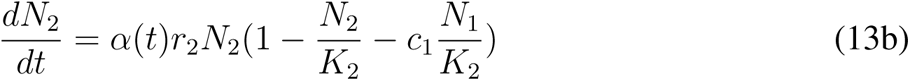

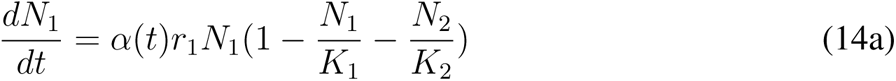

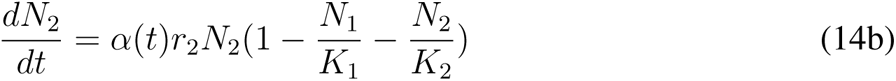

The relative fitness is calculated using:

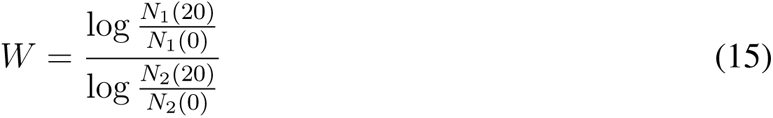

**Figure 7:**
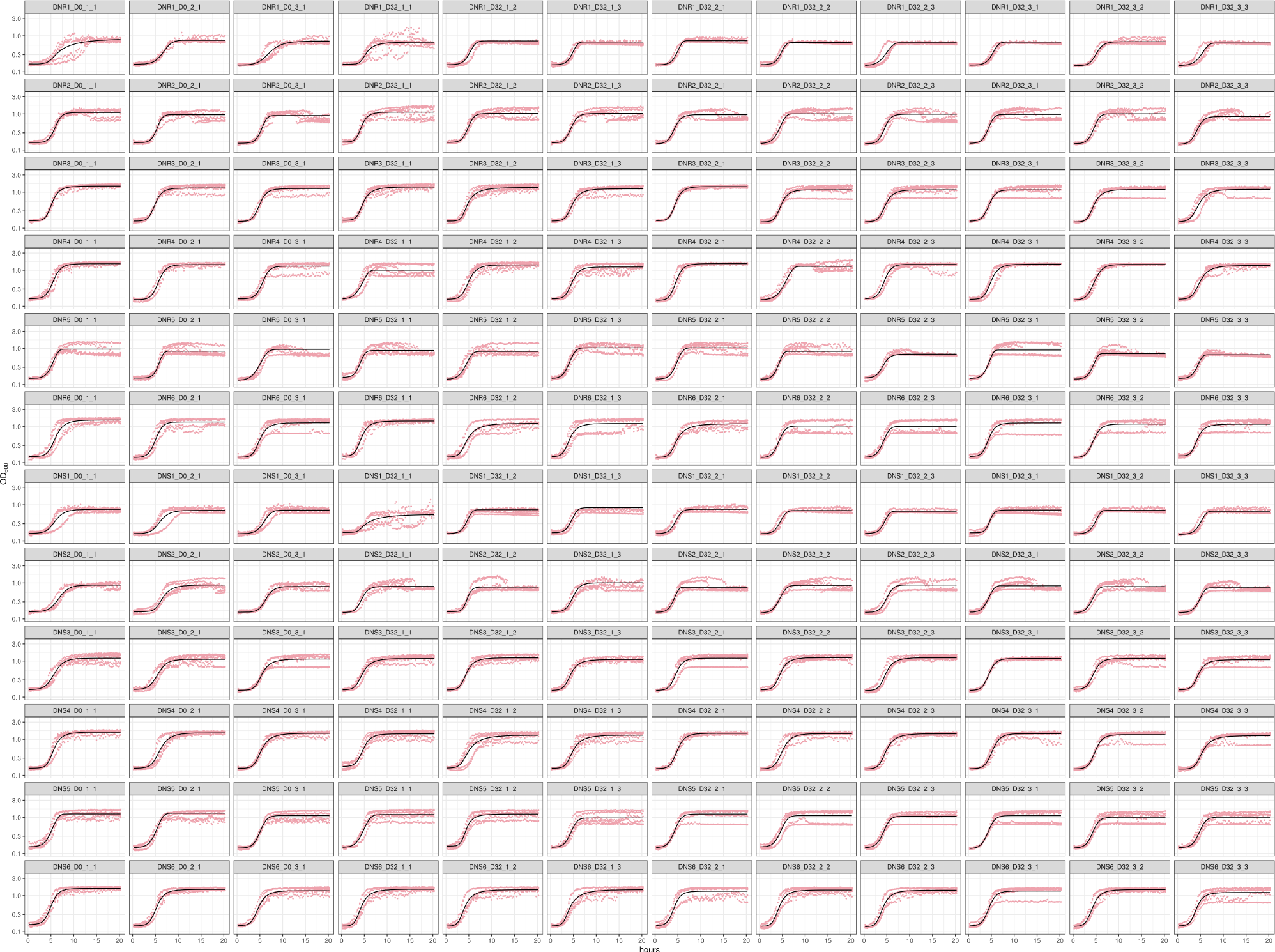
Growth curves of *de novo* resistant strains. Each point represents one of 2 technical replicates from one of the three different biological replicates. The best fit from the growth model is superimposed. Each plot is labeled by Strain ID, Day, Experimental Population, and Clone

**Figure 8:**
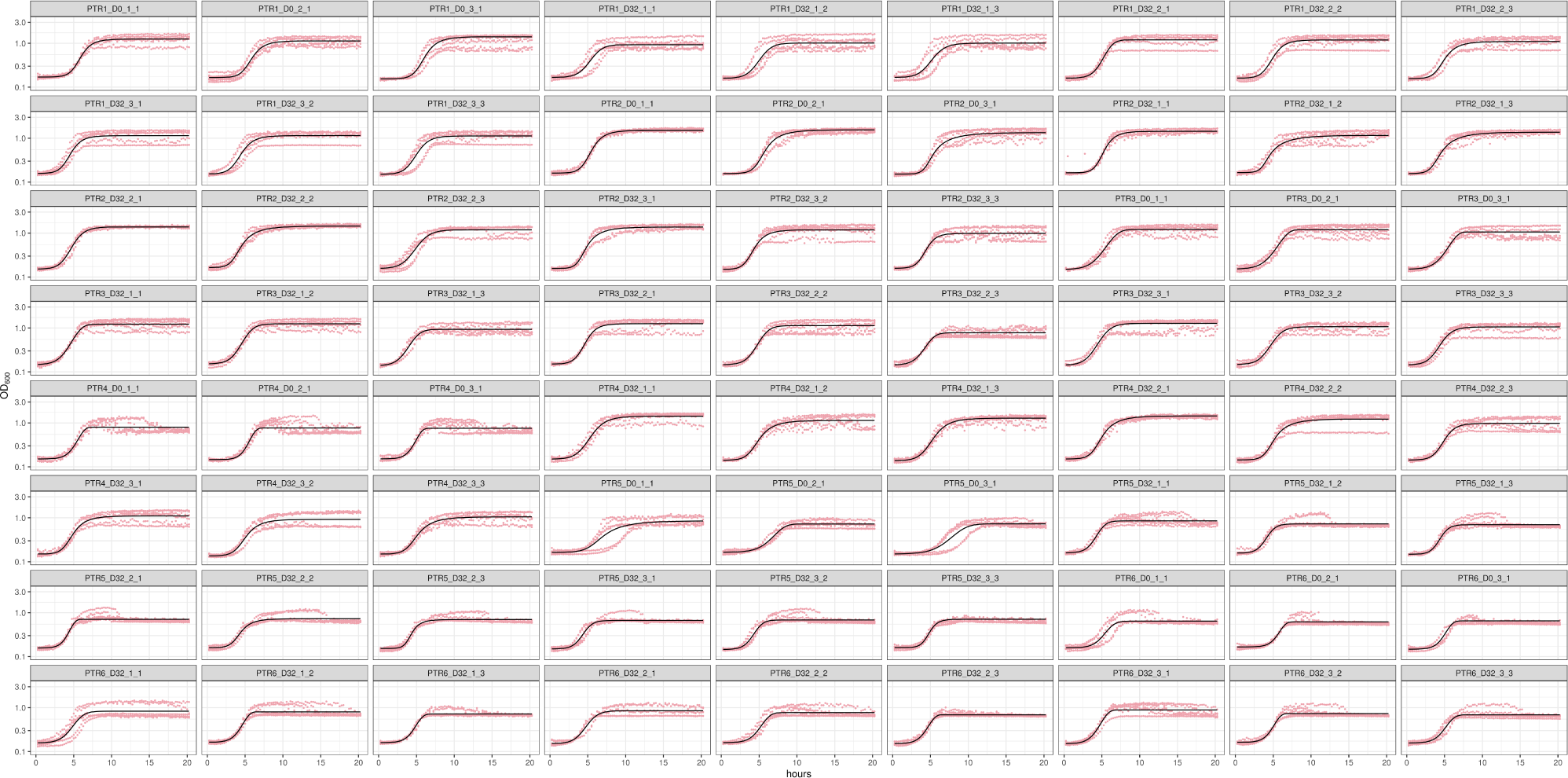
Growth curves of transmitted resistant strains. Each point represents one of 2 technical replicates from one of the three biological replicates. The best fit from the model is superimposed. Each plot is labeled by Strain ID, Day, Experimental Population, and Clone

**Figure 9:**
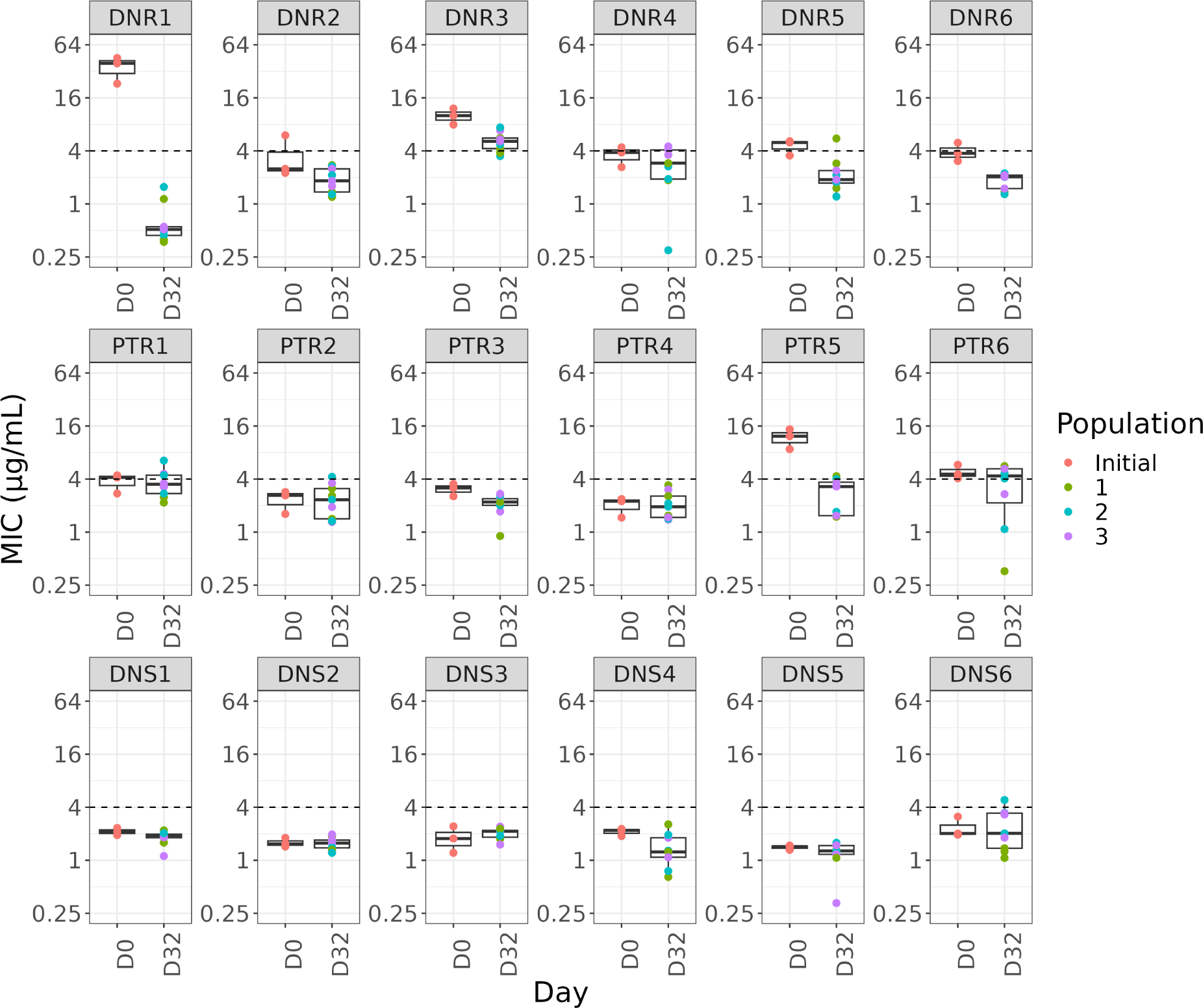
Daptomycin resistance levels before and after evolution in an antibiotic-free environment. Minimum inhibitory concentration of daptomycin from clinical isolates (initial) and three isolates from each of three replicate populations following 320 generations of experimental evolution.

